# The reproductive microbiome inhibits pollen germination in milkweed

**DOI:** 10.1101/2025.09.25.678039

**Authors:** Harmony J. Dalgleish, Joshua R. Puzey, Katie Barlow, Olivia Cunningham, Elizabeth M. Davies, Hannah Machiorlete, Geneva Waynick, Kurt E. Williamson

## Abstract

We know very little about the reproductive microbiomes of plants. Microbes may play important roles in shaping pollination, fertilization, and seed production – processes which are important evolutionarily, ecologically, and agriculturally. Through a series of field and laboratory experiments, we show that the stigmatic microbiome in milkweeds influences the success of pollination. Isolation of individual bacterial and fungal taxa from stigmatic secretions allowed us to experimentally test their effects on pollen germination. These experiments demonstrate that individual taxa impact pollen differently, with many microbial taxa being neutral, but some being deleterious. Through isolation of microbes from the legs of pollinator insects we found that pollinators are a likely source for pollen-harmful bacterial taxa. Next, by utilizing a natural hybrid zone, we demonstrate species-specific responses to the stigmatic microbiome that be driving asymmetric patterns of gene-flow between species – with *Asclepias exaltata* being a better pollen host than *A. syriaca*. This study demonstrates that the reproductive microbiome is an underappreciated player in sexual reproduction of plants.

**Significance Statement:** The results presented here demonstrate an important but previously unappreciated role of stigmatic microbes in plant sexual reproduction. This study demonstrates that the microbial taxa living in stigmatic secretions in milkweed impact pollen germination. We found that filtering out the microbes from stigmatic secretions of milkweed flowers dramatically increases pollen germination. Through isolating microbial taxa from both stigmatic secretions, and pollinator legs, we found that individual microbial taxa impact pollen differently, with many taxa being neutral, but some being deleterious. Finally, by utilizing a naturally occurring milkweed hybrid zone we demonstrated that microbial taxa in stigmatic secretions may be acting as asymmetric prezygotic barrier.

## Introduction

The explosion of research into the microbes that live in and on our bodies (the microbiome) has revolutionized our understanding of human health (1, 2). While an abundance of studies have focused on the gut and oral microbiomes, only recently has the reproductive microbiome been understood as contributing to sexual health and fertility, beyond the study of sexually transmitted infections (3). For example, the vaginal microbiota of mammalian females has been connected to fertility and pregnancy outcomes, including in humans (4–6) and agriculturally relevant mammals (7), particularly cattle (8–10). The interaction between male and female reproductive microbiomes may drive fertility, hybrid incompatibility and may influence the evolutionary ecology of all sexually reproducing organisms (3). Despite the expansion of research into the reproductive microbiomes of animals, we know very little about the reproductive microbiomes of plants. Microbes may play a role in shaping pollination, fertilization, and seed production – processes which are important evolutionarily, ecologically, and agriculturally.

Milkweeds are an excellent model system to understand the impact of the reproductive microbiome in plants because of their unique pollination system (Fig 1a). Like many crop plants (e.g., *Malus, Prunus, Vaccinium, Medicago*), milkweed has a wet pollination system which requires moisture on the stigma for pollen tubes to germinate and grow long enough to transport the sperm to fertilize the ovule (11, 12). In milkweed, thousands of pollen grains are packaged into a structure called pollinia that can be seen with the naked eye and easily manipulated (Fig 1a). Moreover, the stigmatic nectar required for pollen germination (13) is abundant and can be easily sampled, described, and manipulated (Fig 1b). Other wet pollination systems rely on sticky fluid on stigmas, often of low volume, making their study more difficult (but see (14)). Also, adding interest to the milkweed system is that pollinia are transported from flower to flower on insect legs, making insects a potential source of microbes to the plant reproductive microbiome. Finally, we have identified a hybrid zone between two species of milkweed, *Asclepias exaltata* and *Asclepias syriaca*, enabling the study of reproductive microbiomes in driving hybridization and patterns of gene-flow.

**Figure 1.**
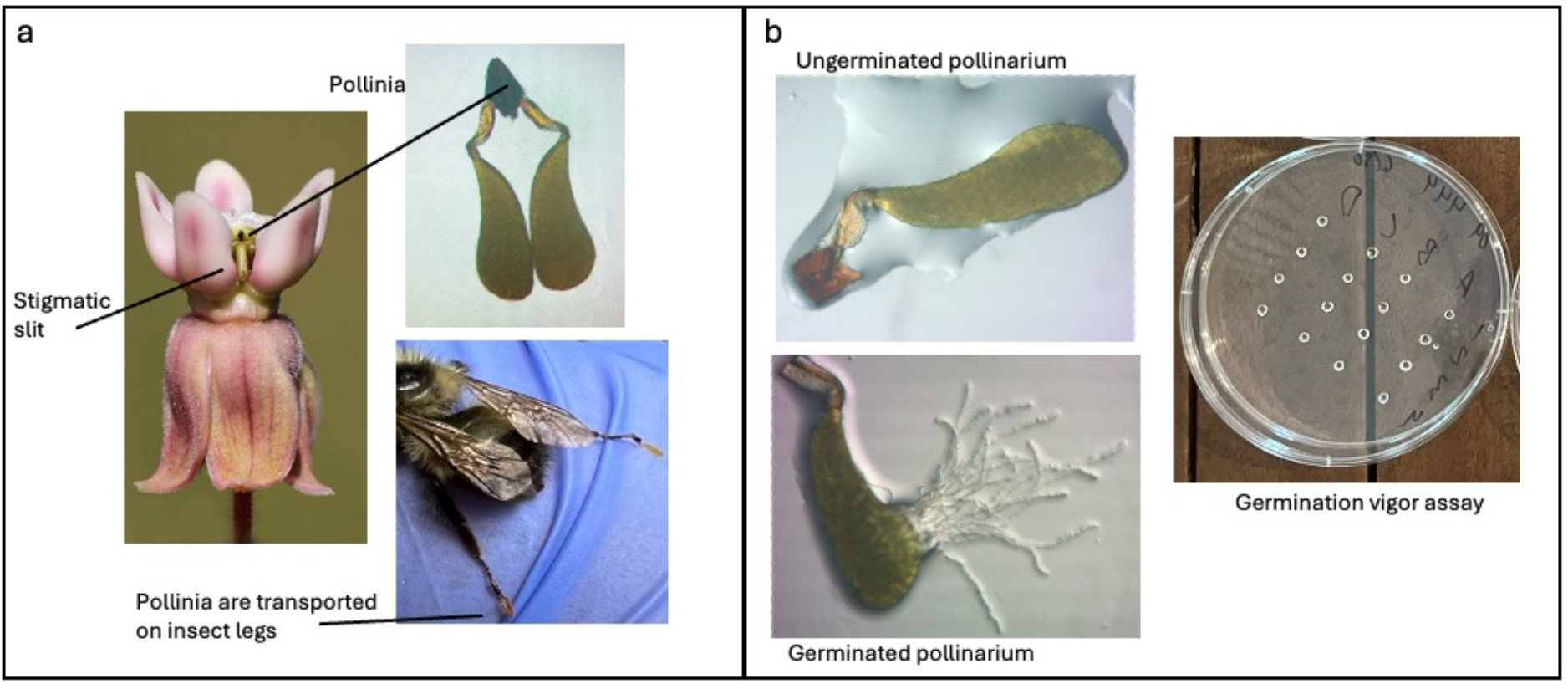
Milkweed floral structure and example of experimental assays. (a) All milkweeds package their pollen into pollinia. Pollinia are transported between flowers on insect legs and inserted into the stigmatic slit, which is filled with nectar, the stigmatic secretion that contains the reproductive microbiome. We isolated 32 bacterial and 31 fungal taxa from insect pollinators and from nectar of *Asclepias syriaca*. (b) Pollinia germination vigor assays consist of drops of nectar (field-collected or artificial) placed on a petri dish, each with a single pollinarium. After ∼8 hours, they are imaged under magnification. Germination vigor is calculated as the sum of z-scores for germination (yes or no), number pollen tubes, and length of longest pollen tube.

In this current study, we have conducted a series of field and laboratory studies to understand the influence of the reproductive microbiome in milkweed on pollen tube germination and growth. Specifically, we have removed the entire reproductive microbiome through filtering, experimentally tested the effects of 63 unique reproductive microbial taxa on pollinia germination, tested the role of pollinators as vectors of harmful microbial taxa, and investigated the role of the reproductive microbiome in driving gene-flow through reciprocal pollen germination experiments in an active hybrid zone between *A. syriaca* and *A. exaltata*.

## Results

### The reproductive microbiome decreases pollen germination vigor in Asclepias syriaca

When we removed microbes through filtering nectar collected from the field at the end of the flowering season in 2023, we found that pollinia germination vigor was higher in filtered nectar (Suppl. Fig. 1, n = 6 nectar collections pooled across multiple inflorescences, *F*_1, 401.9_ = 90.64, *P* < 0.0001; using both nectar collection pool and pollinia source as random effects). We repeated this experiment in 2024, sampling nectar throughout the entire flowering season (n = 12 pooled nectar collections). Our results remained consistent between years: pollinia germination vigor increased when microbes were removed through filtering (Fig 2a). In addition, germination vigor in the filtered samples decreased from the early to the late season (Fig 2a, treatment * season: *F*_*2, 1001.87*_ = 245.08, *P* < 0.0001; using both nectar collection pool nested within season and pollinia source as random effects). Germination in unfiltered nectar remained consistently low across the season (Fig 2a). Even in the late season, however, pollinia germination vigor was higher when microbes were removed through filtering (*P* < 0.001).

**Figure 2.**
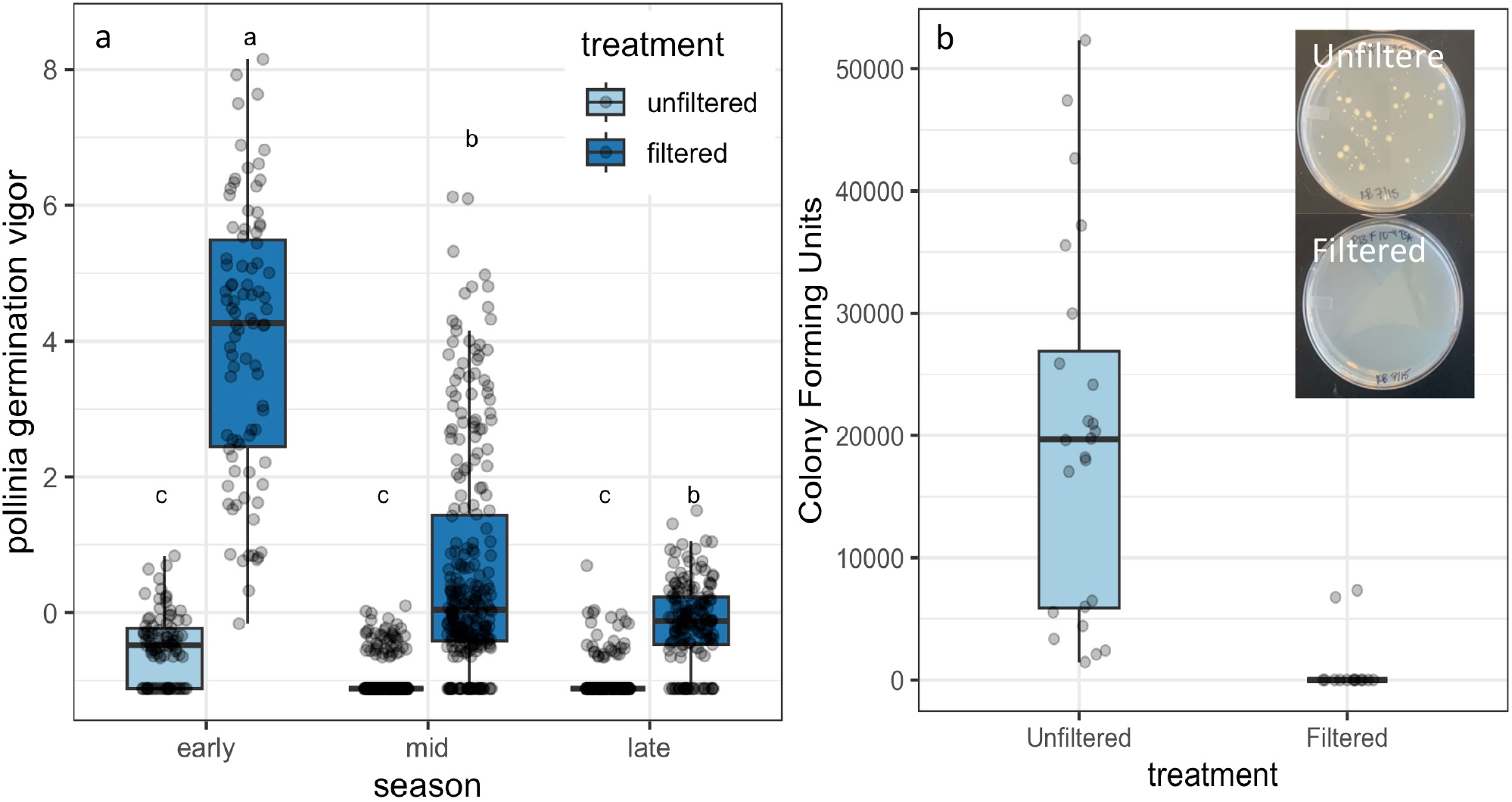
Effects of filtering to remove the microbiome from whole nectar. (a) Germination vigor of pollinia increased when field-collected nectar of *Asclepias syriaca* was filtered to remove the reproductive microbiome. Germination vigor also decreased across the season from early June (early) to July (late) (treatment * season: F_2, 1001.87_ = 245.08, P < 0.0001; using both nectar collection nested within season and pollinia source as random effects) b) Unfiltered nectar had a mean of 20,082 CFUs when plated on TSA agar. Filtered nectar had drastically reduced CFUs with mean of 1006 CFUs and all but two pools having no CFUs when plated on TSA agar. *F*_*1, 36*_ *= 22.34, P < 0.0001*.

The filter treatment dramatically reduced the average number of colony forming units (CFUs) from ∼20,000 uL^-1^ to ∼1000 uL^-1^ (Fig 2b, *F*_*1,36*_ *=* 22.34; *P* < 0.0001). The majority of filter treatments (80% of the 12 pools) reduced CFUs to zero.

### Pollinator vectored microbiomes negatively impact pollen germination

Preventing pollinator access through bagging virgin inflorescences significantly increased the average and variance of pollinia germination vigor (Fig 3a). When we individually tested the effects of 63 distinct microbial taxa isolated either from whole nectar or cultured from common milkweed pollinators in artificial nectar, we found 30% of them were antagonistic, *i.e*., they significantly reduced pollinia germination vigor below pure artificial nectar controls (Suppl. Fig 2). In addition, antagonistic microbes were more likely to be cultured from insects than from nectar (Fig 3b, *χ*^*2*^_*Pearson*_ *= 16.15, P < 0.0001*, n = 63) and were more likely to be bacterial than fungal (Fig 3c, *χ*^*2*^_*Pearson*_ *= 12.09, P < 0.0001*, n = 63). Flowers that were accessible to pollinators had reproductive microbiomes dominated by bacteria compared to bagged flowers (Fig 3d, 67% amplified bacteria only vs. 33% that amplified only fungi or both bacteria and fungi, *P < 0.0001*). In addition, the reproductive microbiome of bagged flowers had lower richness of the common bacterial taxa (Fig 3e).

**Figure 3.**
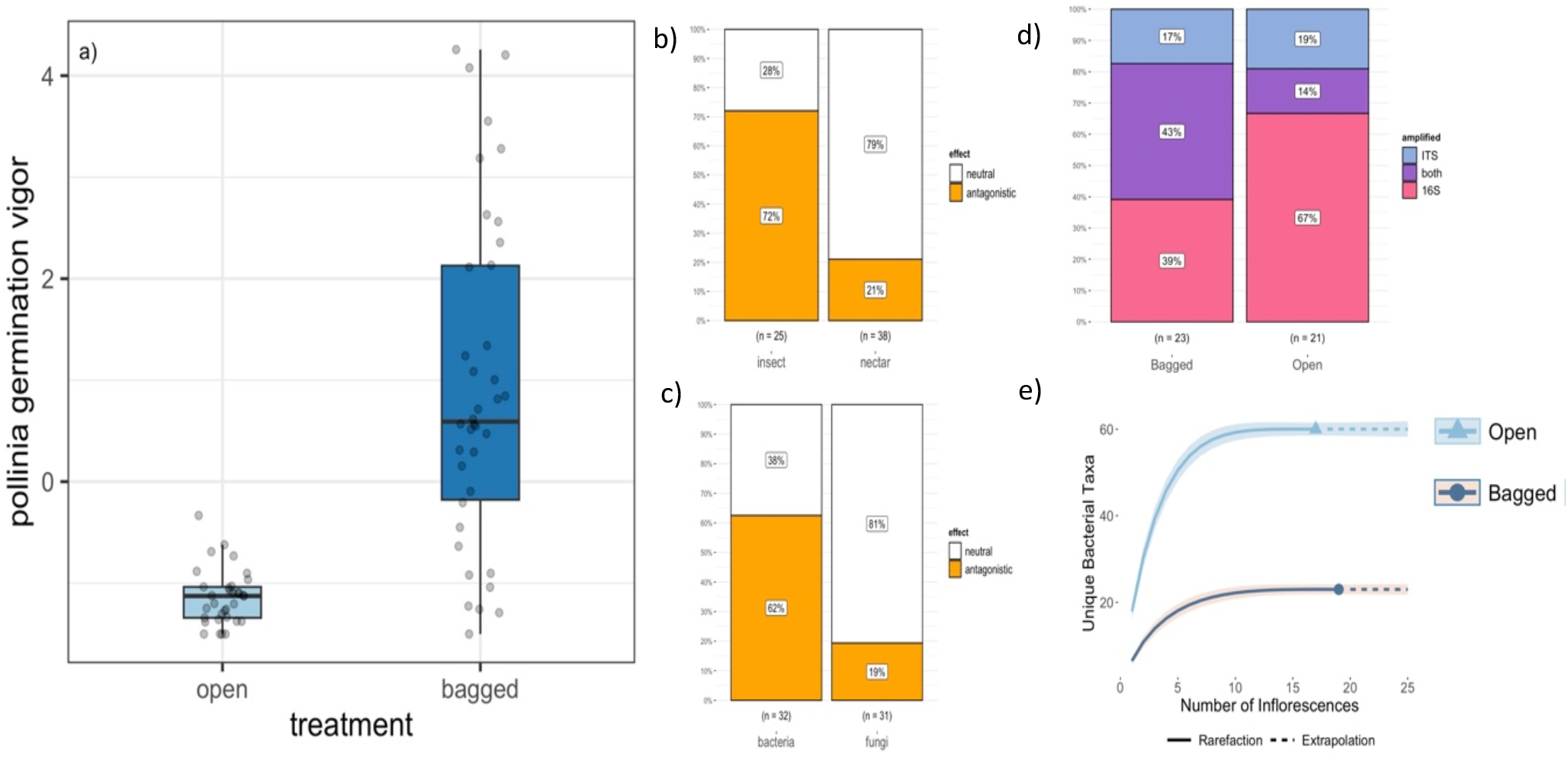
Effects of excluding pollinators on microbiome composition and pollinia germation. (a) Pollinia germination vigor decreased in inflorescences open to pollinators for 24 hours. Inflorescences that were bagged had higher and more variable pollinia germination vigor. (b) Microbial taxa isolated from insects (n = 25) were more likely to be antagonistic to pollinia germination (either prevent or significantly decrease germination vigor) than microbial taxa isolated from nectar (n = 38), *P < 0.0001*. (c) Regardless of sources, bacteria (n = 32) are more likely to be antagonistic to pollinia germination vigor than fungi (n = 31), *P < 0.0001*. (d). Open inflorescences were more likely to have reproductive microbiomes dominated by bacteria than reproductive microbiomes dominated by fungi (67% vs 33%, *P < 0.0001*). e) Reproductive microbiomes in open inflorescences had a higher number of common bacteria taxa than bagged inflorescences. Shaded area is 95% confidence interval.

### The reproductive microbiome alters hybridization by creating asymmetric pollen germination

*Asclepias syriaca* and *A. exaltata* co-occur and readily hybridize along the Appalachian Mountains in the eastern US (15, 16). *Ascepias exaltata* occurs in the shady forest understory and *A. syriaca* occurs in sunny, open areas such as power line cuts. Hybrids have been found along forest edges. In addition to habitat preferences, the leaf shape, presence of trichomes on the underside of the leaves, flower number per inflorescence and flower color differentiate *A. syriaca, A. exaltata* and their hybrids (Fig. 4a). Whole genome resequencing revealed bidirectional introgression between these species (Fig. 4b; Suppl Table 1 lists sampling locations). We have found hybridization to be asymmetric: *A. exaltata* individuals are generally more admixed than *A. syriaca* (Figure 4b and Suppl Fig 3). This pattern is not easily explained by differential abundance as *A. exaltata* is numerically more abundant at this site than *A. syriaca*.

**Figure 4.**
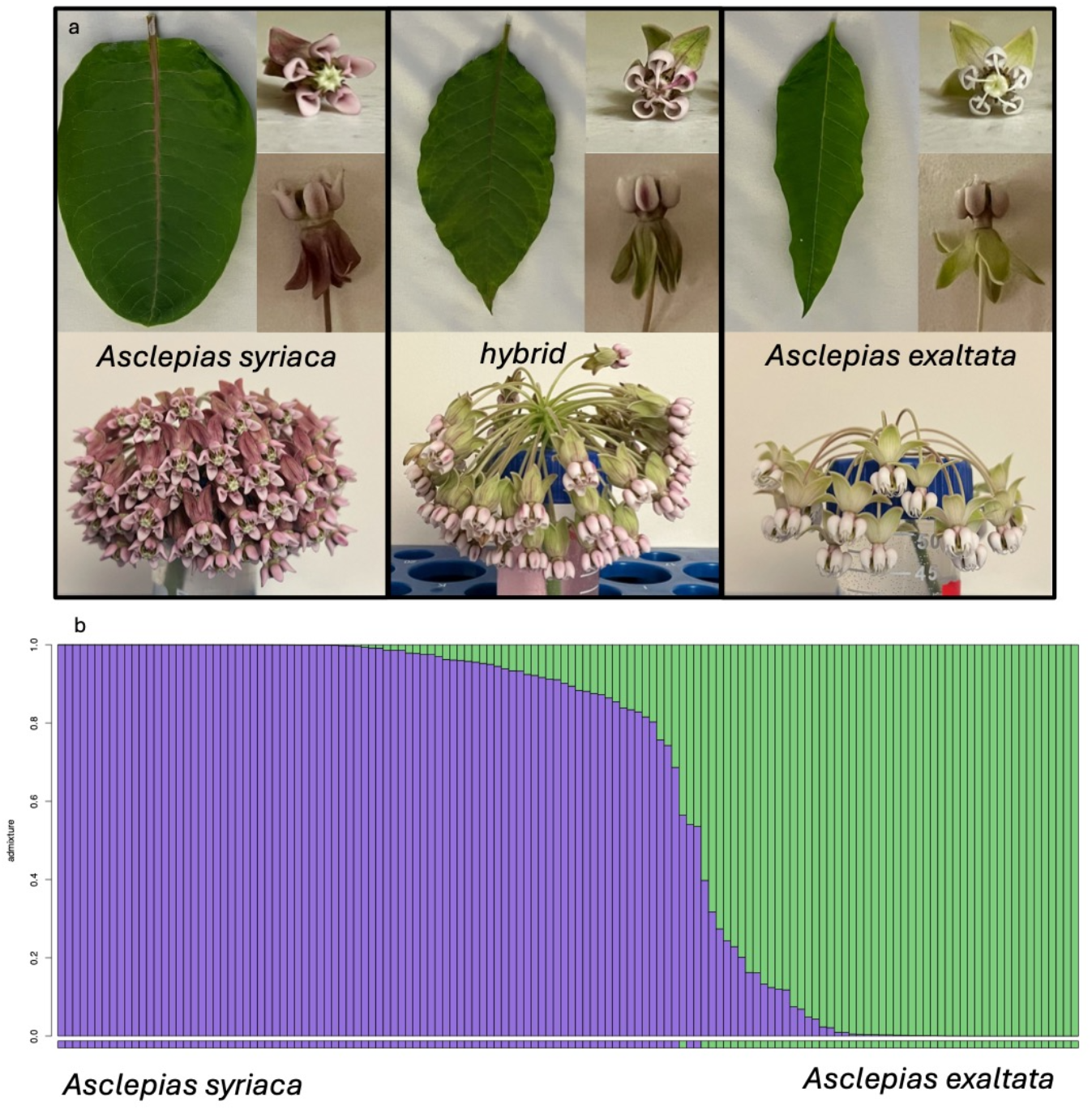
Phenotypic and genotypic characteristics of two milkweed species. (a) Several phenotypes differentiate *Asclepias syriaca, A. exaltata*, and their hybrids including leaf shape, presence of trichomes on the underside of the leaves, flower number per inflorescence and flower color. (b) Hybridization is asymmetric in this population genes from *A. syriaca* are more likely to be present within individuals, even those with *A. exaltata* phenotypes.

Pollinia from both *A. syriaca* and *A. exaltata* had significantly higher germination vigor in nectar collected from *A. exaltata* than nectar collected from *A. syriaca* (Fig 5a, Suppl Fig 4). Further, the reproductive microbiome was quite different between species. Using barcode sequencing of ITS and 16S regions, we found that the reproductive microbiome of *A. syriaca* contained bacteria and fungi, while only fungi were detected in the reproductive microbiome of *A. exaltata* (Fig. 5b). In addition, the density of microbes in the *A. exaltata* reproductive microbiome was significantly lower for *A. exaltata* than for *A. syriaca* (Fig. 5c, Suppl Fig 5). The lower density of microbes and absence of bacteria in the reproductive microbiome of *A. exaltata* enhances pollinia germination of *A. syriaca* within *A. exaltata* compared to germination in *A. syriaca*.

**Figure 5.**
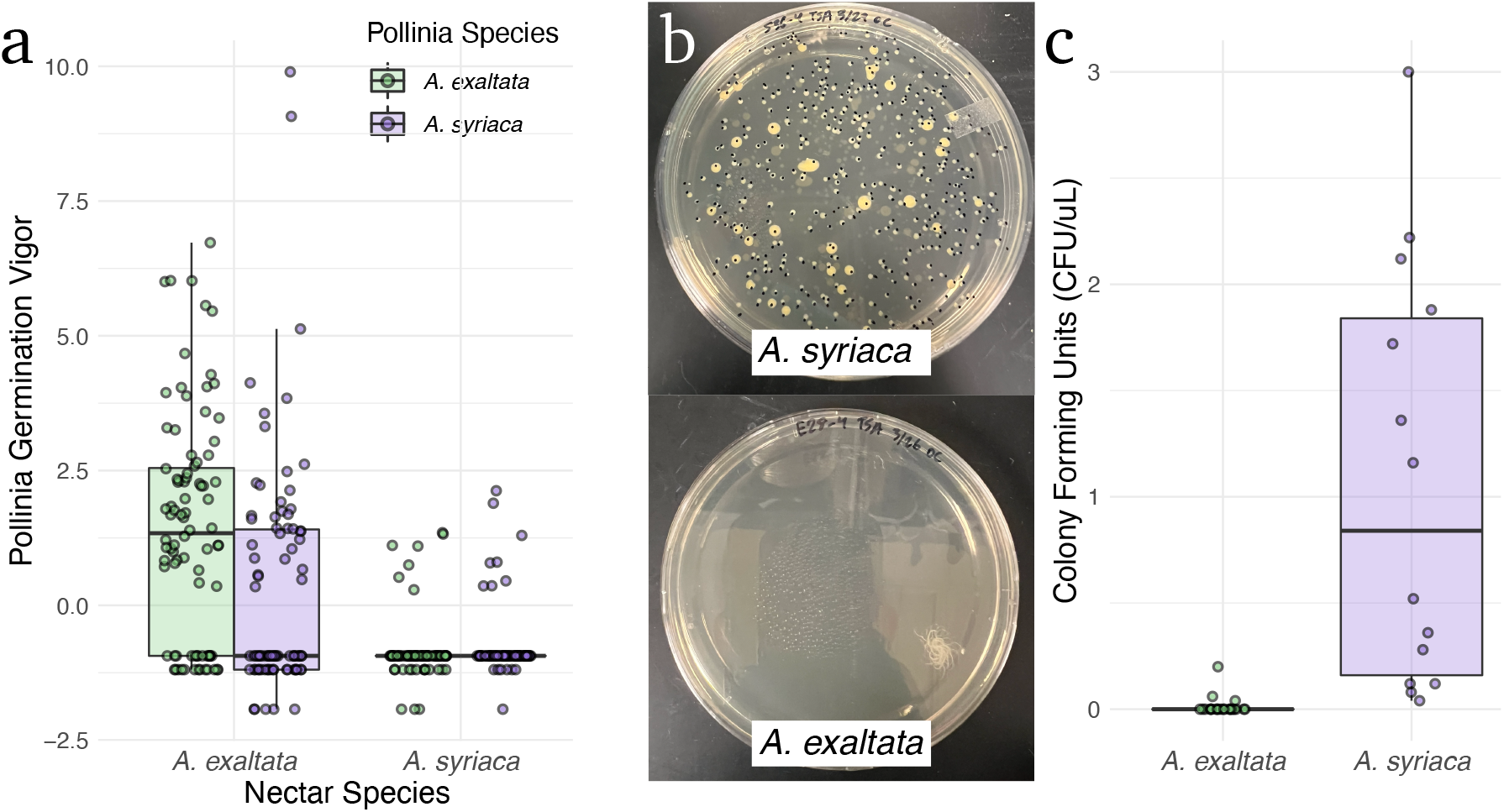
Effects of conspecific nectar on pollinia germination. (a) Pollinia germination vigor was the highest in nectar from *A. exaltata*. b) Nectar plated from *A. exaltata* resulted in low or no colony growth. Nectar plated from *A. syriaca* resulted in high colony growth. c) Nectar plated from *A. exaltata* had significantly fewer CFUs.

## Discussion

Taken together these results demonstrate that the reproductive microbiome of milkweed influences the success of pollination and can be a strong prezygotic barrier. Many angiosperms have stigmatic secretions that facilitate pollen germination (17), and most gymnosperms produce pollination drops, chemically similar to nectar and also consumed by insects, as a germination medium (18). Our work demonstrates that the reproductive microbiome contained within secretions that facilitate pollen tube germination could be an important part of plant sexual reproduction broadly. Within angiosperms, the reproductive microbiome may be an underappreciated third player in the plant-pollinator mutualism, sometimes functioning as a venereal disease for plants vectored by pollinators that can directly prevent successful fertilization and, thus, seed production.

Despite early work in common milkweed showing that some strains of nectar yeast (*Metschnikowia* spp) may inhibit pollen tube germination (19), we found that bacterial taxa are more likely to be antagonistic than fungal taxa. In other plant systems, bacteria have been shown to change pollinator behavior, indirectly decreasing seed production (20). For example, bumblebees (*Bombus* spp.), honeybees (*Apis mellifera*), and mosquitos (*Culex pipiens*) have been shown to prefer nectar containing yeast over bacteria (21, 22). In addition, nectar-associated bacteria have been shown to reduce pollinator attraction in *Mimulus aurantiacus*, which led to reduced pollination success and seed set (23). In an artificial nectar solution, the common nectar bacterium *Acinetobacter* was shown to induce pollen bursting in *Eschscholzia californica* pollen, which would prevent normal tube formation and successful fertilization (24). Our results support that bacteria may be more detrimental to plant reproduction than yeasts, although the mechanisms remain to be explored.

The discovery of a natural hybrid zone between two species of milkweed, *A. exaltata* and *A. syriaca* provided a unique opportunity to study the role of stigmatic secretions in plant hybridization. Interestingly, this hybrid system appears to be asymmetric, with more *A. syriaca* alleles moving into *A. exaltata*, than vice-a-versa. This is surprising given the local dominance of *A. exaltata:* based on plant abundance, we would have predicted that *A. exaltata* alleles would be swamping out *A. syriaca* alleles. Our experimental work with the reproductive microbiome of *Asclepias* demonstrates that *A. exaltata* is a significantly better pollen host than *A. syriaca*. Moreover, the microbiota of the stigmatic secretions differs between these plant species, with *A. exaltata* having only yeast, and *A. syriaca* having both yeast and bacteria. We’ve also observed that bacteria taxa are more antagonistic to pollen germination than fungal taxa in milkweed. These results could explain our observation of asymmetric hybridization between *A. syriaca* and *A. exaltata*, suggesting that the reproductive microbiome may alter evolution by changing gene-flow patterns in plants.

Our results, combined with the recent literature on the important role of reproductive microbiomes in mammals, illustrate the need for expanded understanding of the role of the reproductive microbiome in plants. Many questions remain, such as how is the plant reproductive microbiome formed? Is it determined through priority effects, as has been shown in nectar microbiomes (25, 26)? While we have found many antagonistic and neutral microbial taxa, are there mutualists? Perhaps there are microbial taxa that are not directly beneficial, but rather beneficial through competing with and excluding antagonists within the community (26, 27)? These and many other questions will help us understand the ecological and evolutionary impact of microbes on plant reproduction and open the door to applications that may enhance the reproduction of agricultural crops.

## Materials and Methods

### Pollinia germination vigor assay

To assay pollinia vigor, we used the ‘hanging drop method’ modified from (13). Inside the lid of a petri dish, we placed 5 uL droplets of the treatment liquid (either pure sugar solutions or field-collected nectar). Within each droplet, we placed a single pollinarium, closed the dish and filled the bottom with a 20% sucrose and water solution to prevent the treatment drops from evaporating. The dish sat on the laboratory bench for 6 hours (sugar solutions) or 24 hours (field-collected nectar), after which the remaining liquid in the treatment drop was wicked up with filter paper. Each pollinarium was then visualized under a light microscope at 100x magnification and imaged with a scale bar for reference.

We collected data from each image using ImageJ (28). We recorded whether each pollinarium germinated and, for those that germinated, we counted the number of pollen tubes and measured the length of the longest tube. Within an experiment, we converted each of these three measures of germination into z-scores and summed the z-score for all three measures to quantify germination vigor.

To ensure that we were using viable pollinia for our germination assay, we tested inflorescences the day prior to the experiment. For each of four inflorescences from four different ramets located > 5 m apart from each other, we obtained 5 pollinia from two flowers per inflorescence. We let them germinate in water (2023) or artificial nectar solution (2024; 7% sucrose solution plus trace amino acids) for 6 hours. If fewer than 4/5 pollinia germinated, that inflorescence was not used.

### Germination vigor of pollinia in hetero- and conspecific nectar

We collected pollinia and nectar (see supplemental methods)from pure *A. exaltata*, pure *A. syriaca* and their hybrids from three locations within or around Wintergreen Resort near Nellysford, VA. We collected *A. exaltata* from Ravens Roost (37.932359, -78.950148), pure *A. syriaca* at Stoney Creek (37.919883, -78.913771), and hybrid from Fortunes Ridge (37.904956, - 78.956076). The hybrid is most likely a single, multi-ramet clone. Because our true sample size of hybrids is only a single clone, the data are not representative of a hybrid population. Despite this, we conducted the full factorial experiment using *A. syriaca, A. exaltata*, and this hybrid and present the results in Suppl Fig 3. However, we base our conclusions for this study only on the data from *A. syriaca* and *A. exlatata*, because we have sufficient replication of genets for these species.

Each pollinia germination assay plate contained three replicate drops of nectar from each of the three species. We randomly selected a flower from each species, crossing pollinia species with nectar sources with every combination of nectar and pollinia on each plate. We replicated this plate design 88 times for a sample size of 792 pollinia from a total of 23 *A. exaltata*, 11 hybrid, and 17 *A. syriaca* nectar samples. We collected nectar over several days from the same inflorescence and pooled the collected nectar into a single sample. We used an ANOVA with Tukey post-hoc test on the z-scores of pollinia germination vigor to test for differences among species in germination vigor. All analyses were conducted in R (29).

### Filtering whole nectar to remove the microbiome

We used nectar from inflorescences located at Blandy Experimental Farm near Boyce, VA (39.066190, -78064356). We collected nectar over the course of 2-3 days to create several separate ∼700 uL pools of nectar. Each pool was a combination of nectar collected from a single inflorescence on 4 - 20 different ramets. Nectar pools were stored at -80×C until they were used for the experiment.

For each experiment, we filtered half of the nectar within each pool using a microcentrifuge tube filter (Corning® Costar® Spin-X® centrifuge tube filters, product number CLS8160). We then set up 6 germination vigor assay plates using pollinia from the three different inflorescences. Each plate contained 8 filtered droplets, 8 unfiltered droplets and 2 control droplets on each plate (DI water or artificial nectar) for a total of 48 pollinia per treatment per pool. In 2023, we had 6 pools for a total of 648 pollinia across both treatments. In 2024, we had 12 pools using 1,404 pollinia. We analyzed the z-scores of pollinia germination vigor using a mixed effects ANOVA with filter treatment as a fixed effect and pool and pollinia donor inflorescence as random effects implemented with *lmer* in R. We conducted post-hoc tests using the *emmeans* function.

In 2024, we plated the filtered and unfiltered treatments on agar to count colony forming units (CFUs) to verify that microbial abundance was decreased using the filtering treatment. We plated both treatments on Tryptic Soy Agar (Carolina Biological Supply, Burlington, NC, Item No. 788421) media and Glucose Yeast Extract media (recipe per liter). We prepared a serial dilution and plated 10^–2^, 10^-3^ and 10^–4^ dilutions. After 24 hours growth on the bench (ca. 22°C), we counted CFUs and converted the dilution to undiluted CFUs uL^-1^. Counts were similar on both media, so we took an average of both as an estimate of microbial abundance. CFU data was analyzed using an ANOVA in R.

### Excluding pollinators to alter the nectar microbiome

Prior to the onset of flowering at Blandy Experimental Farm, we randomly chose 33 ramets in the field, ensuring that each was > 1 m from the nearest experimental ramet. When an inflorescence appeared close to opening, we bagged the inflorescence. Bags remained in place and were checked daily until 50%-75% of the flowers on the inflorescence had opened when the ramet was randomly assigned one of two treatments. For the open treatment, bags were removed in the morning and replaced in the evening (∼10 hours later); for the closed treatment, the bag remained in place the entire treatment. The morning after the treatments were in place, we collected nectar. We sampled a total of 71 inflorescences from 33 ramets, with 33 inflorescences in the open treatment and 38 in the closed treatment.

For each inflorescence, we set up three replicate pollinia germination assay petri dishes per treated inflorescence using three nectar droplets per assay and pollinia from three different pollinia donor ramets for a total of 9 pollinia assays per treated inflorescence. We analyzed the z-scores of pollinia germination vigor using a mixed effects ANOVA with bag treatment (open for 24 hours or continually bagged) as a fixed effect and pollinia donor inflorescence as a random effect implemented with *lmer* in R.

### DNA extraction

DNA was extracted from each nectar sample from bagged and open inflorescences using the DNEasy Blood & Tissue kit (Qiagen, Redwood City, CA) according to manufacturer instructions. The Zymo Clean and Concentrate (Zymo, Irving, CA) kit was used to clean and concentrate extracted DNA. Cleaned and concentrated DNA was stored at -20ºC until sequencing.

### 16S and ITS amplicon sequencing

We sent a minimum volume of 10 µl cleaned and concentrated DNA from each nectar sample and four constructed communities (see supplement) to Novogene (Beijing, China) for 16S and ITS amplicon sequencing using the Illumina NovaSeq 6000 platform (San Diego, CA). For each sample, 5 µl of DNA was used for each 16S and ITS sequencing runs. The 16SV4 region was targeted using primers 515F: 5’-GTGCCAGCMGCCGCGGTAA-3’ and 806R: 5’-GGACTACHVGGGTWTCTAAT-3’ to produce 250 - 300 bp reads. The ITS1-5F region was targeted using primers ITS5-1737F: 5’-GGAAGTAAAAGTCGTAACAAGG-3’ and ITS2-2043R: 5’-GCTGCGTTCTTCATCGATGC-3’ to produce 50 - 350 bp reads.

### Sequence processing

Sequences were processed using the dada2 (30) pipeline in R. Novogene supplied demultiplexed reads with adapters and primers removed. For 16S samples, forward and reverse reads were truncated to 200 bp length, with a maximum expected error rate of 3. For ITS samples, forward and reverse reads were filtered to a minimum length of 50 bp and a maximum expected error rate of 2.

### Microbial community data analysis

ASV read counts were normalized to relative abundance within a sample to account for different library sizes. After agglomerating, normalizing, and pruning the ASV table, bagged and open samples were split into separate ASV tables for subsequent analysis. In accordance with the constructed community results (see supplement), we pruned all bacterial ASVs that comprised less than 0.13% of reads combined across all umbels within a treatment. Similarly, we pruned all fungal ASVs that comprised less than 0.2% of reads combined across all umbels within a treatment and did not prune based on frequency.

### Testing the effects of cultivable isolates on pollinia germination

We created a culture library of cultivable microbes from both field-collected nectar and from frequent insect visitors to common milkweed (see supplement). We tested 63 isolates over 8 days, with 8 isolates tested per day using pollinia germination assay for each isolate. We collected six inflorescences per day from which 12 plates were made. Each inflorescence produced 2 plates and each plate consisted of pollinia from only one inflorescence. Each isolate was tested on 9 pollinia. Three different inflorescences donated 3 pollinia each to this total to account for inter-inflorescence variability.

### Preparation of isolate cultures

Two days preceding a pollinia germination assay, each of the 8 isolates for that day were transferred into 5 ml liquid general media (TSB for bacteria and GYE for fungi) and incubated at 27ºC for 24 hours. Then we transferred 100 µl of culture into 5 ml artificial nectar. The artificial nectar included 7.5% sucrose, 3 mM amino acids (sodium caseinate), and was filter sterilized using 0.2 µm, 50 mm diameter vacuum filter (Fisher Scientific, Waltham, MA). We incubated the artificial nectar cultures along with uninoculated artificial nectar as a control at 27ºC for 24 hours. Pollinia germination experiments were conducted with 24-hour artificial nectar cultures.

### Identifying microbes detrimental to pollinia germination

To test if isolates effected pollinia germination vigor, we used a Bayesian linear regression model in the brms package in R (31). The best performing model according to leave one out comparison used a student’s t distribution and normally distributed priors (0, 10) for the regression coefficient, gamma distributed (2, 0.1) for the shape of the student distribution (nu), and exponential distribution (0.1) for sigma. The model was run on 4 chains and cores with 4000 iterations with 2000 warmup iterations. We modeled vigor scores as a function of isolate with the random effects of day, umbel, and plate accounted for as a varying nested intercepts. We set the reference level to the control group.

### Genetic analysis of Ascelpias syriaca and Asclepias exaltata

We visited 16 sites in the mountains of Virginia to collect *A. syriaca* and *A. exaltata* (Table S1). For each plant collected, we measured phenotypic characteristics of each plant and collected two leaves for DNA extractions. Within a single site, we tried to ensure individuals were as far apart as possible to minimize sampling clones. We attempted to sample plants at least 1m or more apart, though this was not always possible. Upon collection, leaves were kept on ice and then frozen at -80×C until we were ready to perform extractions.

We extracted DNA from each milkweed sample using Qiagen DNeasy Plant Mini Kits according to manufacturer instructions. We sequenced the DNA using low-pass whole genome Illumina sequencing at Michigan State University yielding 150bp-PE reads.

We aligned DNA sequences to the milkweed genome using bowtie2 and variants were called using GATK Haplotype Caller (32, 33). We filtered the resultant VCF file using vcftools and the following criteria to ensure high-quality SNPs analyses [--minDP 2 --maf 0.1 --max-missing 0.8 - -min-alleles 2 --max-alleles 2]. Next, SNPs were filtered for linkage using Plink (-indep-pairwise 50 5 0.2) (34). An initial PCA using this filtered revealed a split between *A. syriaca* and *A. exaltata* along PC1. PC2 split *A. syriaca* into one large cluster, and a smaller sub-cluster. The small sub-cluster was solely from one isolated population on the top of Cole Mountain, VA. Also, the two *A. syriaca* individuals that fall within the *A. exaltata* cluster are also from Cole Mountain (Suppl Fig 6). We removed all Cole Mountain samples from subsequent analyses as we were primarily interested in *A. syriaca* and *A. exaltata* population differentiation, not finer aspects of *A. syriaca* population structure. After filtering out the CMB individuals, we retained a total of 138 individuals.

Using the original VCF, except with CMB individuals removed, we filtered using the same criteria as above. The average sequencing depth after this filtering step yielded 4777 SNPs with an average depth of 13.8x.

Next, we used conStruct to determine population structure (35). We ran a non-spatial model for 20,000 iterations at k=2. We chose this K-value because the PCA (with Cole Mountain removed) clearly shows two clusters. The ConStruct analysis revealed a clear signal of introgression between these two species. Using the resultant estimates, we calculated the percent ancestry in hybrid individuals as an estimate of introgression. For individuals with mixed ancestry (those <=97% pure), *A. exaltata* showed different patterns of genetic admixture than *A. syriaca* (Supp Fig 3; Kolmogorov-Smirnov test, *P* = 0.038). A Kolmogorov-Smirnov test was chosen as the assumptions for a t-test were not met.

## Supplementary Information

is available for this paper and provided as a PDF.

## Acknowledgments

We gratefully acknowledge the staff at Blandy Experimental Farms, The Nature Foundation at Wintergreen, Ivan Munkres, Nicki Gustafson, Hether Natterer, and Mia Perry for field assistance as well as the following funding sources:

William & Mary, Arts & Sciences Faculty Research Seed Grant (HJD, JRP, KEW)

Garden Club of America (KB, OC, HM, GW)

Garden Club of Virginia Graduate Fellowship (GW)

Blandy Experimental Farms Graduate Research Fellowship (HM, GW)

W&M Plumeri Award (JRP)

