## Supplemental Information for "The reproductive microbiome inhibits pollen germination in milkweed"

*Harmony J. Dalgleish

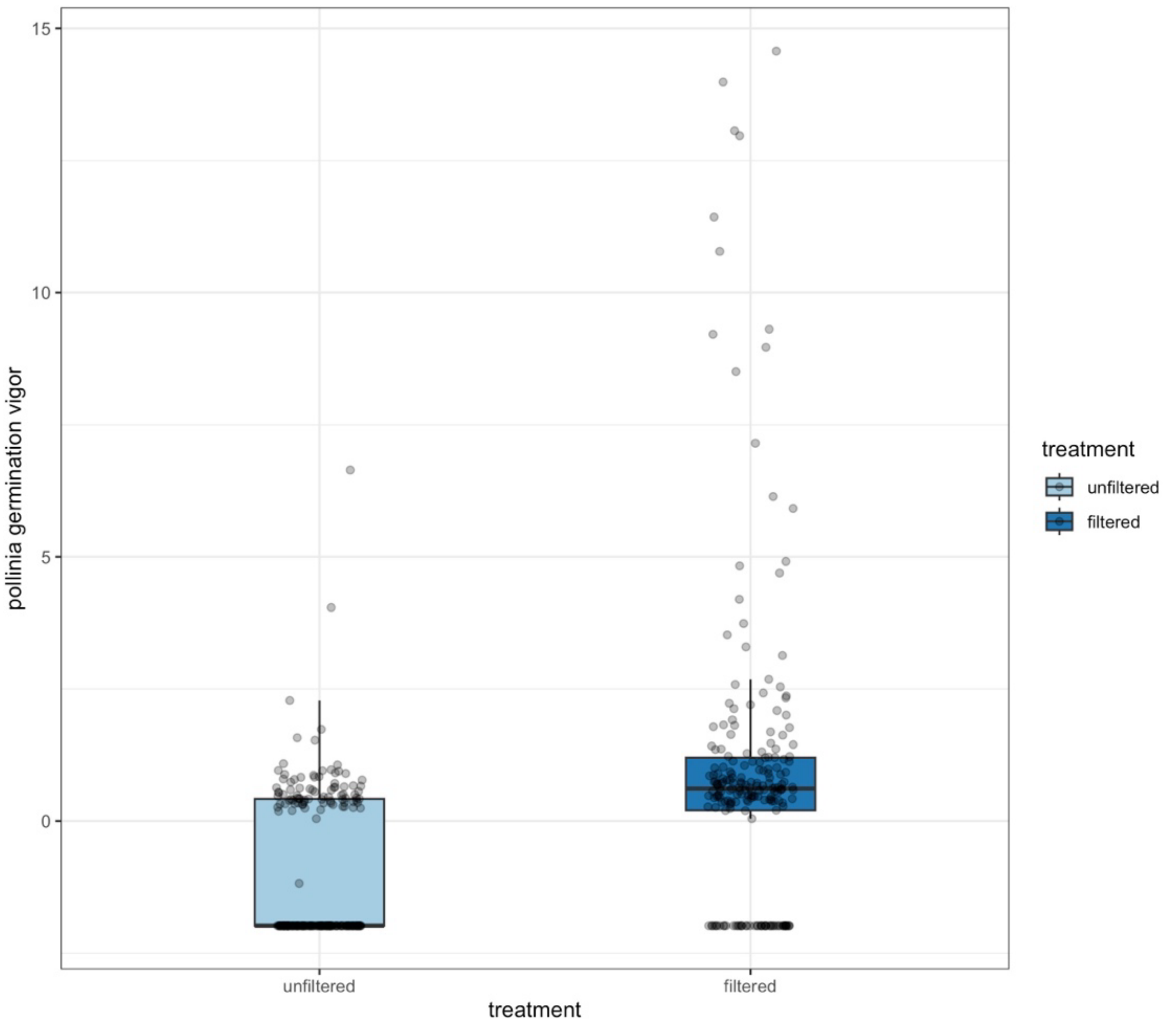


**Fig. S1.** Germination vigor of pollinia increased when field-collected nectar of *Asclepias syriaca* was filtered to remove the reproductive microbiome. n = 6 nectar collections pooled across multiple inflorescences, $F_{1, 401.9} = 90.64$, $P < 0.0001$; using both nectar collection pool and pollinia source as random effects. Data from 2023. When we repeated the experiment in 2024, we found the same results (see Fig. 2a in the main text).

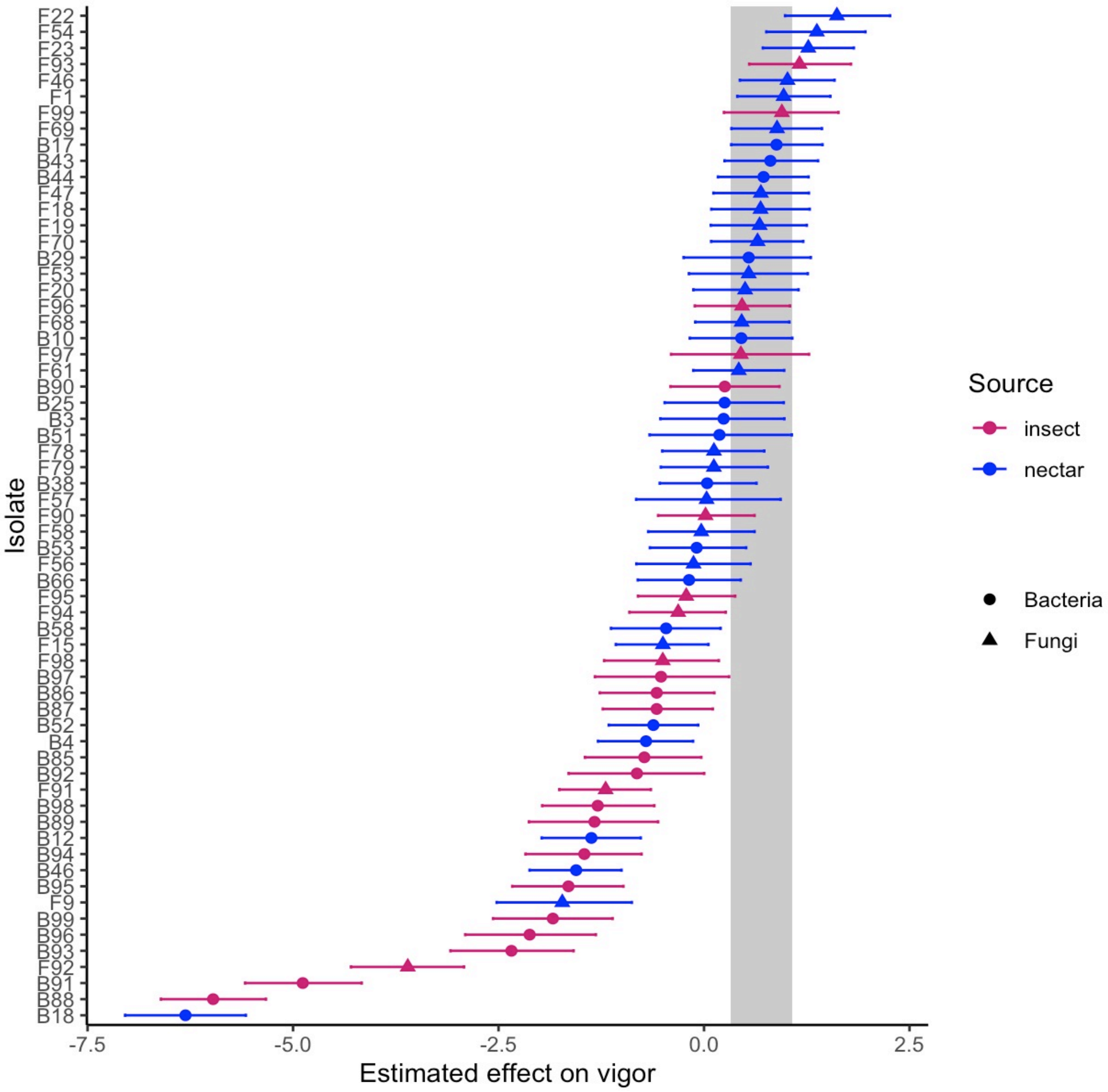


**Fig. S2.** Twenty-six of 63 isolates decreased pollinia germination compared to control (41%). No isolate increased pollinia germination. Each isolate is colored by source (insect or nectar) with the different symbols representing bacteria (circle) or fungi (triangle). Grey bar represents the 80% credible interval of the control (pollinia germination in artificial nectar).

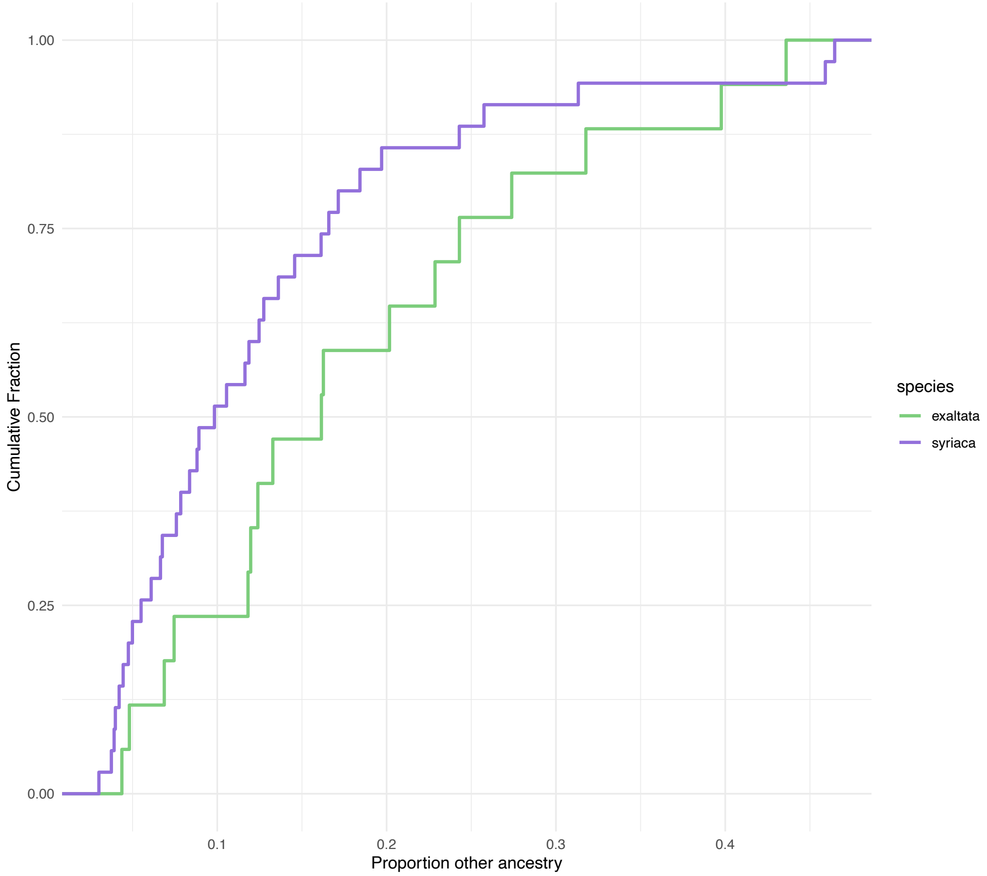


**Fig. S3.** Cumulative Distribution Plot of Genetic Admixture. *A. exaltata* generally has higher levels genetic admixture based on ancestry estimates from ConStruct (Kolmogorov-Smirnov test, $P = 0.038$).

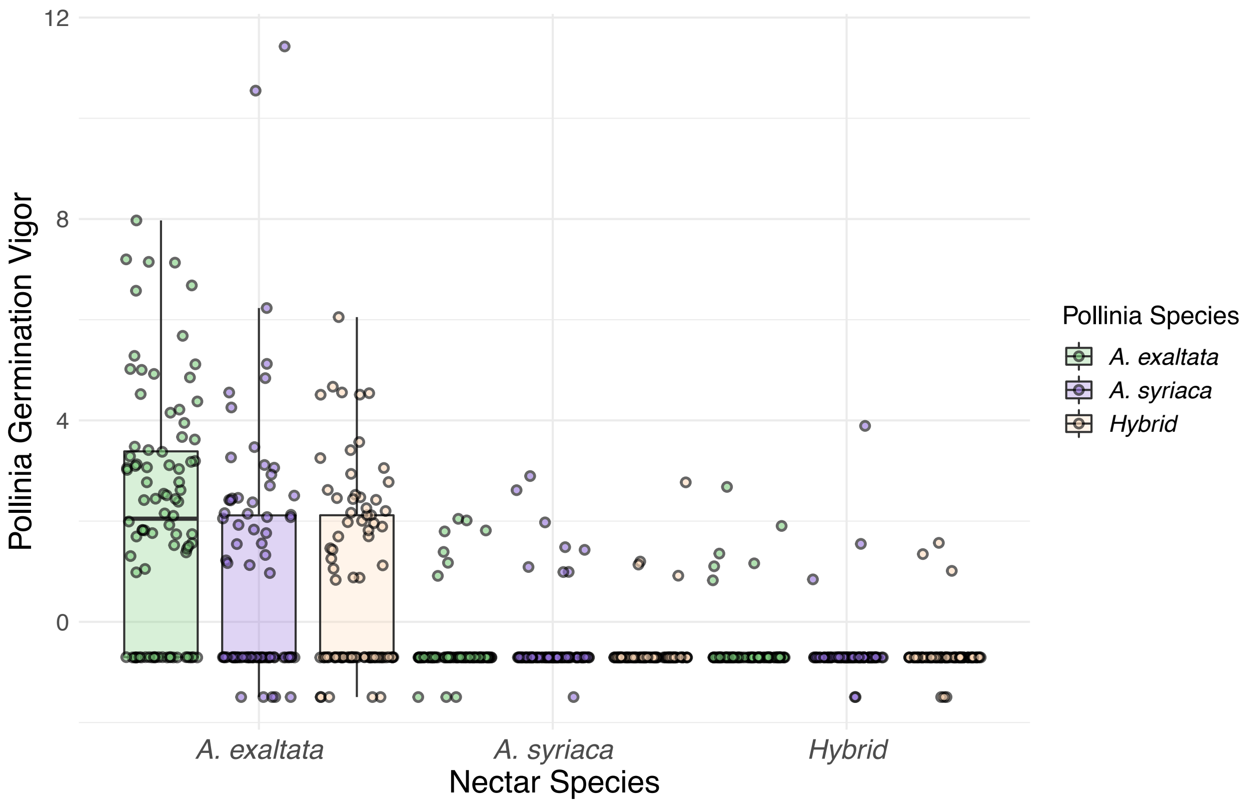


**Fig. S4.** Samples were largely split by species along the PC1 axis. One population of *A. syriaca* samples existing alone on the top of a single mountain (Cole Mountain, labeled in orange) formed a separate cluster along the PC2 axis. Points are labeled based on their phenotypic call in the field

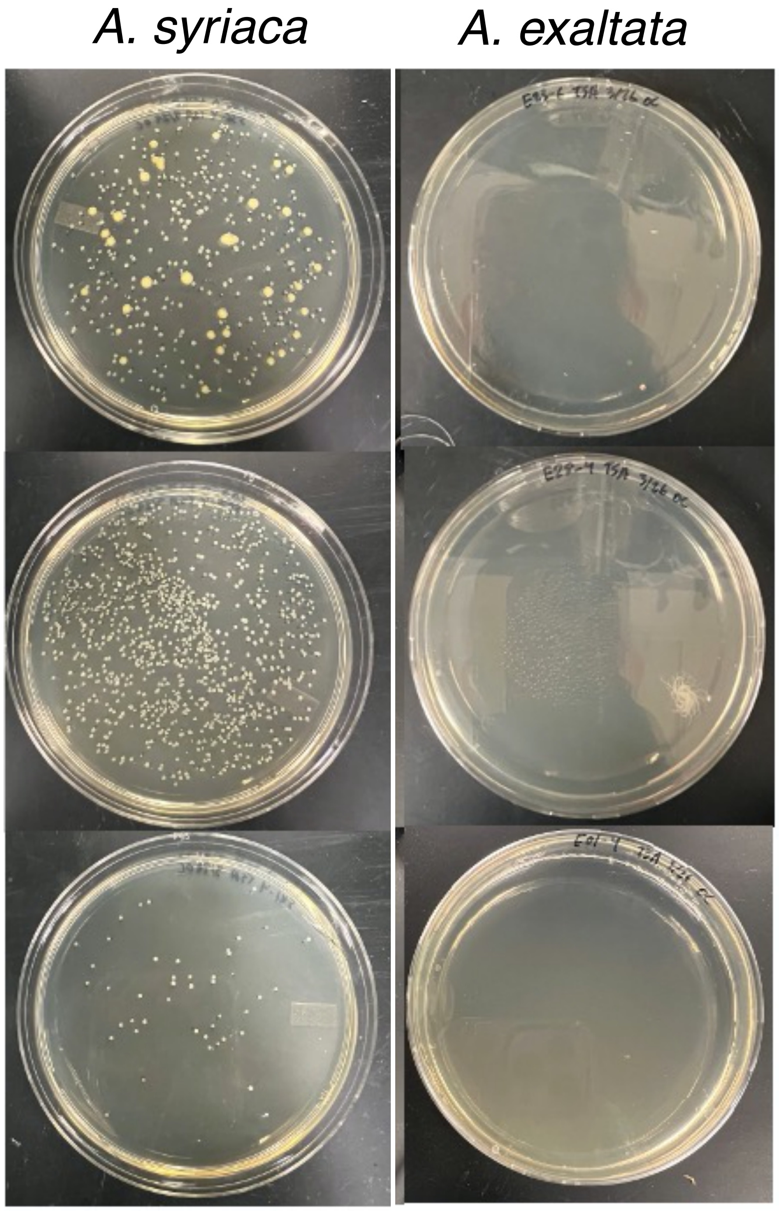


**Fig. S5.** Representative samples of plates from the CFU experiment consistently show *A. syriaca* with larger numbers of CFUs than *A. exaltata*

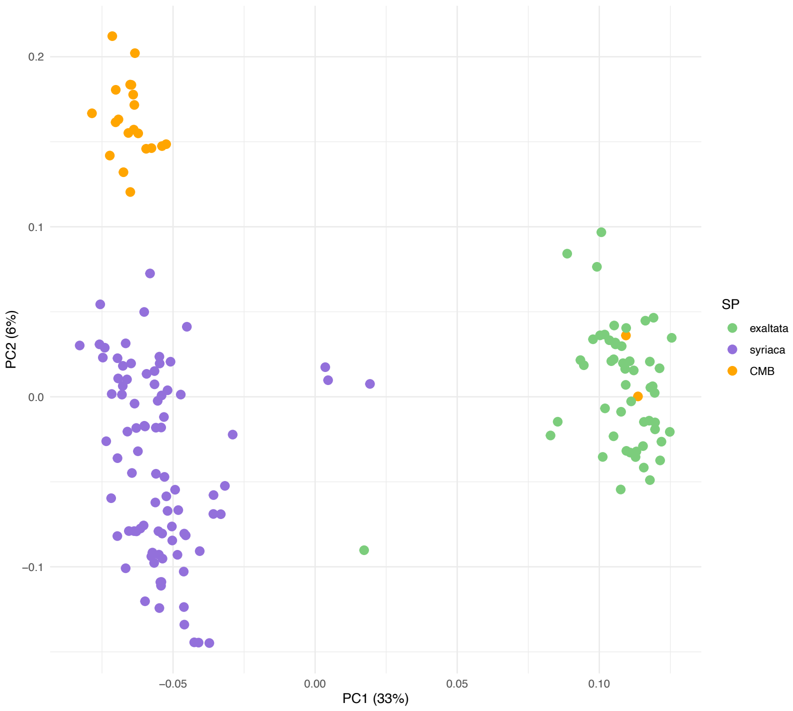


**Fig. S6.** Samples were largely split by species along the PC1 axis. One population of *A. syriaca* samples existing alone on the top of a single mountain (Cole Mountain, labeled in orange) formed a separate cluster along the PC2 axis. Points are labeled based on their phenotypic call in the field.

| Site Key | Full Site Name | Latitude | Longitude | # *A. syriaca* collected | # *A. exaltata* collected |
|---|---|---|---|---|---|
| LM | Lower Mashokin | N 37° 54.944’ | W 078° 54.635’ | 20 | 0 |
| HR | Humpback Rocks | N 37° 56.6826 | W 78° 54.6882 | 0 | 0 |
| GH | Glass Hollow | N 37° 56.6826 | W 78° 54.6882 | 0 | 0 |
| RGT | Rockfield Gap Tourist Site | N 38° 01.827’ | W 078° 51.799’ | 7 | 0 |
| PTW | Peddar Trail Wintergreen | N 37° 54.579’ | W 78° 55.918’ | 0 | 10 |
| FRW | Fortunes Road Wintergreen | N 37° 54.288’ | W 078° 57.355’ | 18 | 0 |
| SLG | Salt Log Gap | N 37° 46.807’ | W 079° 10.924’ | 15 | 7 |
| LFS | Lush Forest Shire | N 37° 46.378’ | W 079° 11.042’ | 0 | 15 |
| MMP | Milkweed Mountain Peak | N 37° 46.017’ | W 079° 11.226’ | 17 | 0 |
| CMB | Cole Mountain Bald | N 37° 45.051’ | W 079° 12.321’ | 21 | 6 |
| RF | Rice Field (AT) | N 37° 22.603’ | W 080° 45.343’ | 12 | 1 |
| PNR | Pembroke Near Rice | N 37° 21.979’ | W 080° 44.680’ | 6 | 0 |
| MKP | Mcafee Knob Parking lot | N 37° 22.808’ | W 080° 05.395’ | 8 | 0 |
| TCP | Tinker Cliff Parking lot | N 37° 27.446’ | W 080° 01.044’ | 10 | 0 |
| RRL | Raven Roost Lane | N 37° 56.022 | W 78° 56.9748 | 0 | 22 |
| SGC | Swannanoa Golf Course | N 38° 1.1934’ | W 78° 52.758’ | 12 | 0 |

**Table S1.** Site names and locations for Milkweed Collections. GPS coordinates are given in Degrees Decimal Minutes (DDM). The number of *A. syriaca* or *A. exaltata* collected refers to the number of plants recognizably characterized as either species in the field.

**Supplementary Methods**

Species descriptions

*Asclepias syriaca* (common milkweed)*,* is a perennial herb that ranges from southern Canada to Georgia, USA, and from the Atlantic Ocean to the central Great Plains in North America. It flowers indeterminately in large inflorescences that can range in color from light pink to purple. The flowers of common milkweed have tall and peaked hoods with small horns (Fig. 1). The leaves of common milkweed are wide with a rounded tip. The leaves tend to have a purple mid-vein and a soft underside due to an abundance of trichomes. Common milkweed is a largely self-incompatible plant (< 5% self-compatibility) and it reproduces clonally by re-sprouting from adventitious root buds (*19*). It thrives in disturbed habitats in full sun.

*Asclepias exaltata* (poke milkweed), is a perennial herb with a range that largely overlaps that of *A. syriaca* in North America but is most abundant along the Appalachian Mountain range. *Asclepias exaltata* flowers indeterminately in small inflorescences that can range in color from pure white to light pink. The flowers of poke milkweed have small and domed hoods with large horns (Figure 1). The leaves of poke milkweed are thin with a pointed tip and tend to have a white mid-vein. They have a waxy underside due to a lack of trichomes. Like common milkweed, poke milkweed is largely self-incompatible and reproduces clonally by re-sprouting from adventitious root buds (*20*). Unlike common milkweed, poke milkweed thrives in forested habitats and partial shade.

The *Asclepias syriaca x Asclepias exaltata* (intermediate hybrid), is a perennial herb that is found in areas where populations of common milkweed and poke milkweed overlap. The hybrid is primarily found around the edges of mountain balds or powerline clearings. It flowers indeterminately in mid-sized inflorescences that can range in color from white to dark pink. The flowers of the intermediate hybrid have short, rounded, open hoods with long horns (Figure 4a). The leaves of the hybrid are wide with a pointed tip. The leaves tend to have a partially purple mid-vein and a soft underside as they have an intermediate amount of trichomes. The grows and flowers in areas where either common or poke milkweed are found.

Field nectar collection protocol

We bagged the inflorescences the evening before with a clean satin bag (Uline, 5x7 inch white satin bags, product number S-21653) and returned the next morning between 05:00 and 07:00. Using a sterile 10 uL microcapillary tube, we collected as much nectar as possible from the flowers within the inflorescence. Nectar was transported in a cooler in the field and stored at -80$^{o}$C until used for pollinia germination assays.

Constructed community sequencing controls

To quantify sequencing biases, we created a mock community control sample consisting of equimolar concentrations of genomic DNA of 10 bacterial and 10 fungal isolates (Table S1). Eight bacterial and 7 fungal genera were represented in the mock community. Four dilutions of mock community control were sent for 16S and ITS amplicon sequencing: 10, 5, 2.5, and 1 ng $\mu l^{-1}$ of total DNA in molecular grade water, representing the approximate range of DNA yield from field collected nectar.

After agglomerating to genus and transforming ASV counts to proportion per sample, we found 12 bacterial genera in the mock community. Out of the eight bacterial genera used in the mock community samples, we detected 5 successfully . Three of the eight genera we added to the community were not detected: *Enterobacter*, *Pantoea*, and *Erwinia*. All genera except one (*Klebsiella*) with more than 0.13% of a sample's reads (30 reads per sample) were actual mock community members. We detected seven genera that were not actual mock community members, but all except for *Klebsiella* had very low read count. Read counts for bacterial genera are approximately similar across the three highest DNA concentrations, which the lowest concentration did not amplify.

We found 31 fungal genera in the fungal mock community. Of the seven fungal genera used in the mock community samples, we reliably detected five and one, *Gibellulopsis,* was not detected. *Pseudocosmospora* was found in only one sample at a very low read count. Taxa with more than 0.2% of reads (100 reads per sample) are all actual mock community constituents. Read counts for fungal genera are approximately similar across different DNA concentrations.

**Isolating cultivable microbes from nectar and insects**

Insect isolates

To isolate microbes found on insects that commonly pollinate milkweed, we collected insects (n = 26) on milkweed at Blandy Experimental Farm and State Arboretum (Boyce, VA). Specimens were collected over a two-week period between mid- and late-July 2022 and were transferred in a cooler for <4 hours before long term storage at -80ºC. We prepared an artificial nectar model (15% sucrose with 3 mM sodium caseinate to supplement amino acids, (*26*)) and inoculated it with surface microbes from insects. For each culture, we poured a small amount of sterile artificial nectar into a weigh boat, then placed a single specimen in the artificial nectar for 30 seconds. The inoculated artificial nectar was then transferred to a sterile 1.5 ml microcentrifuge tube. Each artificial nectar culture was inoculated with a single specimen, then we incubated the culture at 25ºC for 48 hours. We prepared serial dilutions of $10^{-1}$, $10^{-2}$, 10-$^{4}$, $10^{-6}$, $10^{-8}$ for each sample using filter-sterilized artificial nectar.

Nectar isolates

To isolate microbes found in nectar, we collected nectar from 9 inflorescences randomly selected at Blandy Experimental Farm during spring 2023. We chose inflorescences that had > 75% of flowers open. We bagged the inflorescences the evening before with a clean satin bag (Uline, 5x7 inch white satin bags, product number S-21653) and returned the next morning between 05:00 and 07:00. Using a sterile 10 uL microcapillary tube, we collected as much nectar as possible from the flowers within the inflorescence. We prepared serial dilutions of $10^{-1}$, $10^{-2}$, 10-$^{4}$, $10^{-6}$, $10^{-8}$ for each sample using phosphate buffered saline as diluent.

Isolating pure cultures

We plated 45 µl of each dilution from each nectar or insect sample onto microbial media and evenly distributed across plate using a flame-sterilized glass L-spreader. We use trypticase soy broth (TSA) to culture bacteria supplemented with 0.2% cycloheximide to prevent growth of fungi and glucose yeast agar (GYA) to culture fungi supplemented with 0.2% chloramphenicol to prevent growth of bacteria. We dissolved antimicrobials in 100% ethanol, filter sterilized with a 0.2 µm vacuum filter and then added them after media was autoclaved. We incubated the plates at room temperature (ca. 22.5ºC) and checked them every 24 hours for growth, recording the time of first colony appearance. We distinguished morphotypes of colonies by color, shape, size, and margin. We recorded the number of morphologically distinct colonies per plate and the colony count per morphotype.

We isolated each morphotype observed per plate by re-streaking a single colony onto new media. This process was repeated until a pure culture of each morphotype (henceforth referred to as isolate) was obtained. We considered a culture pure if all colonies on the plate appeared morphologically identical. Once all cultures were determined pure and containing only the targeted isolate, we stored each isolate in 30% glycerol at -80ºC.

DNA extraction

We streaked isolates from frozen glycerol stocks onto the appropriate media type. A single colony was collected from each pure isolate plate and transferred to broth medium and incubated in broth in a shaking incubator set to 200 rpm and 28ºC for 24 hours.

For each isolate, we obtained a cell pellet by spinning 1.5 ml of broth culture in a microcentrifuge at 14,000 rpm for 5 minutes. We aspirated the supernatant and then resuspended the cell pellet in 175 µl of lysis buffer (400 mM NaCl, 750 mM sucrose, 20 mM EDTA, 50 mM Tris-HCl pH 8.0) using a combination of vortexing and pipetting. We added 2 µl of 100 mg $ml^{-1}$ lysozyme (MP Biomedicals, LLC, Solon, OH) and mixed by flicking tube, then incubated in a 37ºC heat block for 30 minutes. We then added 20 µl of 10% SDS and 1 µl 100 mg $ml^{-1}$ proteinase K (New England Biolabs, Ipswich, MA) and mixed by flicking the tube, then incubated in a 55ºC heat block for 1.5 hours. We added 20 µl of 3M sodium acetate and mixed by inverting tube and then added 200 µl of phenol/chloroform/isoamyl alcohol (25:24:2) and mixed by inverting tube several times, then spun at 14,000 rpm in a refrigerated centrifuge set to 4ºC for 5 minutes. We transferred the aqueous layer to a new tube and added 100 µl lysis buffer to bulk aqueous layer, then added 200 µl chloroform and mixed by inverting tube several times, then spun at 14000 rpm in a refrigerated centrifuge set to 4ºC for 5 minutes. We transferred the aqueous layer to a new tube, then added 2 µl Glycoblue (Fisher Scientific, Walham, MA) and 200 µl ice-cold isopropanol and mixed by inverting tube. We obtained a DNA pellet by spinning the tube at 14,000 rpm in a refrigerated centrifuge set to 4ºC for 5 minutes. We aspirated the excess isopropanol and allowed it to evaporate from DNA pellet in a biosafety cabinet. We then resuspended the dried DNA pellet in 25 µl of molecular grade water, then stored at -20ºC.

We used NanoDrop One to measure DNA yields for each isolate (ThermoFisher, Walham, MA) and recorded the DNA concentration (ng $\mu l^{-1}$) and purity (A260/A280; A260/A230). Samples with a minimum of 100 ng $\mu l^{-1}$ were selected for sequencing.

Sanger sequencing

For bacterial isolates, we used primers 16 SF1: 5'-GASTTTGATCCTGGCTYAG-3' and 16 SR1: 5'-GACGGGCGGTGWGTRCA-3' (Integrated DNA Technologies, Coralville, IA) to target the 16S region and produce a ca. 1000 bp sequence length. For fungal isolates, we used primers ITS1 F: 5'-TCCGTAGGTGAACCTGCGG-3' and ITS4 R: 5'-TCCTCCGCTTATTGATATGC-3' (Integrated DNA Technologies, Coralville, IA) to target the ITS region and produce variable sequence lengths.

We used the QIAquick PCR Purification Kit (Qiagen, Hilden, Germany PCR) to purify products and we again measured DNA concentration and purity yields using NanoDropOne (ThermoFisher, Walham, MA). Samples with a minimum of 100 ng $\mu l^{-1}$ purified amplicons were submitted for sequencing.

All 46 insect bacteria samples were sequenced at the University of Delaware Sequencing and Genotyping Center (Newark, DE) and 86 samples (23 insect fungi, 31 nectar fungi, 32 nectar bacteria) were sequenced at the William and Mary Core Lab (Williamsburg, VA) using Sanger technology.

Sequence processing

Sequences were analyzed using the DNASubway [dnasubway.cyverse.org] pipeline (*27*). We used the 'Determine Sequence Relationships' track to assemble sequence pairs and produce a consensus sequence. Using the consensus sequences, we found the closest matching taxonomic identification for each isolate though searching in the BLASTN database.
